# Comprehensive evaluation of *ACE2* expression in female ovary by single-cell RNA-seq analysis

**DOI:** 10.1101/2021.02.23.432460

**Authors:** Siming Kong, Zhiqiang Yan, Peng Yuan, Xixi Liu, Yidong Chen, Ming Yang, Wei Chen, Shi Song, Jie Yan, Liying Yan, Jie Qiao

**Affiliations:** Center for Reproductive Medicine, Department of Obstetrics and Gynecology, Peking University Third Hospital, Beijing 100191, China; National Clinical Research Center for Obstetrics and Gynecology, Beijing 100191, China; Key Laboratory of Assisted Reproduction (Peking University), Ministry of Education, Beijing 100191, China; Beijing Key Laboratory of Reproductive Endocrinology and Assisted Reproductive Technology, Beijing 100191, China; Beijing Advanced Innovation Center for Genomics, Beijing 100871, China; Peking-Tsinghua Center for Life Sciences, Peking University, Beijing 100871, China; Research Units of Comprehensive Diagnosis and Treatment of Oocyte Maturation Arrest; Academy for Advanced Interdisciplinary Studies, Peking University, Beijing 100871, China

**Keywords:** SARS-CoV-2, *ACE2*, primordial germ cells, oocytes, cumulus-oocyte complex

## Abstract

Pneumonia induced by severe acute respiratory coronavirus 2 (SARS-CoV-2) via ACE2 receptor may affect many organ systems like lung, heart and kidney. An autopsy report revealed positive SARS-Cov-2 detection results in ovary, however, the developmental-stage-specific and cell-type-specific risk in fetal primordial germ cells (PGCs) and adult women ovary remained unclear. In this study, we used single-cell RNA-sequencing (scRNA-seq) datasets spanning several developmental stages of ovary including PGCs and cumulus-oocyte complex (COC) to investigate the potential risk of SARS-CoV-2 infection. We found that PGCs and COC exhibited high *ACE2* expression. More importantly, the ratio of *ACE2*-positive cells was sharply up-regulated in primary stage and *ACE2* was expressed in all oocytes and cumulus cells in preovulatory stage, suggesting the possible risk of SARS-CoV-2 infection in follicular development. CatB/L, not TMPRSS2, was identified to prime for SARS-CoV-2 entry in follicle. Our findings provided insights into the potential risk of SARS-CoV-2 infection during folliculogenesis in adulthood and the possible risk in fetal PGCs.

## 1. Introduction

The outbreak of coronavirus disease 2019 caused by SARS-CoV-2 has been reported over 22,000,000 cases of infections. Common symptoms of SARS-CoV-2 infected patients were fever, fatigue, dry cough, anorexia, and myalgia (Wang et al., 2020). The severe acute respiratory syndrome (SARS) infected binding receptor ACE2 was also thought to be the receptor for SARS-CoV-2 as SARS-CoV-2 shared similar sequences in receptor-binding domain with SARS (Wan et al., 2020) and was confirmed to invade cells via ACE2 by *in vitro* experiments (Zhou et al., 2020). Meanwhile, Hoffmann *et al* found that the serine protease TMPRSS2 activity plays a crucial role in SARS-CoV-2 priming and endosomal cysteine proteases CatB/L may also have some effects in TMPRSS2 negative cells (Hoffmann et al., 2020). Therefore, cells with *ACE2* and *TMPRSS2* expression and *TMPRSS2* negative cells with *ACE2* and *CatB/L* expression, may act as target cells and are susceptible to SARS-CoV-2 infection. Previous single-cell RNA-sequencing (scRNA-seq) analysis were mainly focused on the *ACE2* expression level of lung, heart, esophagus, kidney, bladder, and ileum (Zou et al., 2020). An autopsy report showed positive detection in ovary (W, 2020), however, the related details about patients such as specifically detective area, number of detective patients, ages were not included. As human ovary is a complex organ and contains different types of cells, systematical *ACE2* expression analysis is critical in evaluation the risk in female ovary.

A key function of ovary is to generate fertilizable oocytes with full competence for reproduction (Oktem and Oktay, 2008). In female embryos, PGCs can form oocytes after controlled cell division and meiosis (Felici, 2005; Nikolic et al., 2016). In adult women, oocytes and cumulus cells in follicles plays a primary role in folliculogenesis (Zhang et al., 2018).

Different cell types may behave distinct susceptibility to SARS-CoV-2. We used our previous scRNA-seq data on PGCs (Li et al., 2017) and COC (Zhang et al., 2018) to construct a potential risk map of SARS-CoV-2 infection in fetal PGCs and adult women ovary.

## 2. Results

### 2.1 Female PGCs exhibited high *ACE2* expression

Female embryos form PGCs during development (FUJIMOTO et al., 1977; Saitou and Yamaji, 2012) and this process includes mitotic, RA signaling-responsive, meiotic, and oogenesis stages (Saitou and Yamaji, 2012). To evaluate the risk of SARS-CoV-2 infection in PGCs, we calculated developmental-stage-specific *ACE2* expression pattern in PGCs. *ACE2* expression were detected in all four cell types, including mitotic (3% *ACE2*-positive cells), RA signaling-responsive phase (10% *ACE2*-positive cells), meiotic prophase (10% *ACE2*-positive cells), and oogenesis phase (59% *ACE2*-positive cells), with an increasing tendency in PGCs formation (Fig 1A, 1B). Previous study regarded cell types with >1% proportion of *ACE2*-positive cells as high risk of infection of SARS-CoV-2 (Zou et al., 2020), therefore, it suggested that PGCs may be at high risk of infection. To further study the potential risk of *ACE2*-positive PGCs, we compared the synchronization of *ACE2* and *TMPRSS2* expression. The results showed that *TMPRSS2* were nearly negatively expressed in PGCs, which revealed that TMPRSS2 may not be used for cell entry by SARS-CoV-2 in PGCs (Fig 1C). We then evaluated the CatB/L expression in *ACE2* positive PGCs. Interestingly, we found that *CTSB* and *CTSL* displayed coincident expression with *ACE2* (Fig.S1A), indicating the possibility of SARS-CoV-2 priming by CatB/L in PGCs. In addition, somatic cells surrounding PGCs, including endothelial cells, early granulosa, mural granulosa, and late granulosa had no *ACE2* and *TMPRSS2* expression (Fig 1D-F). Although CatB/L was expressed in somatic cells surrounding PGCs (Fig.S1B), these somatic cells were still at low risk for SARS-CoV-2 infection. Taken together, PGCs may be at high risk while their surrounding somatic cells may be at low risk for SARS-CoV-2 infection.

**Figure 1.**
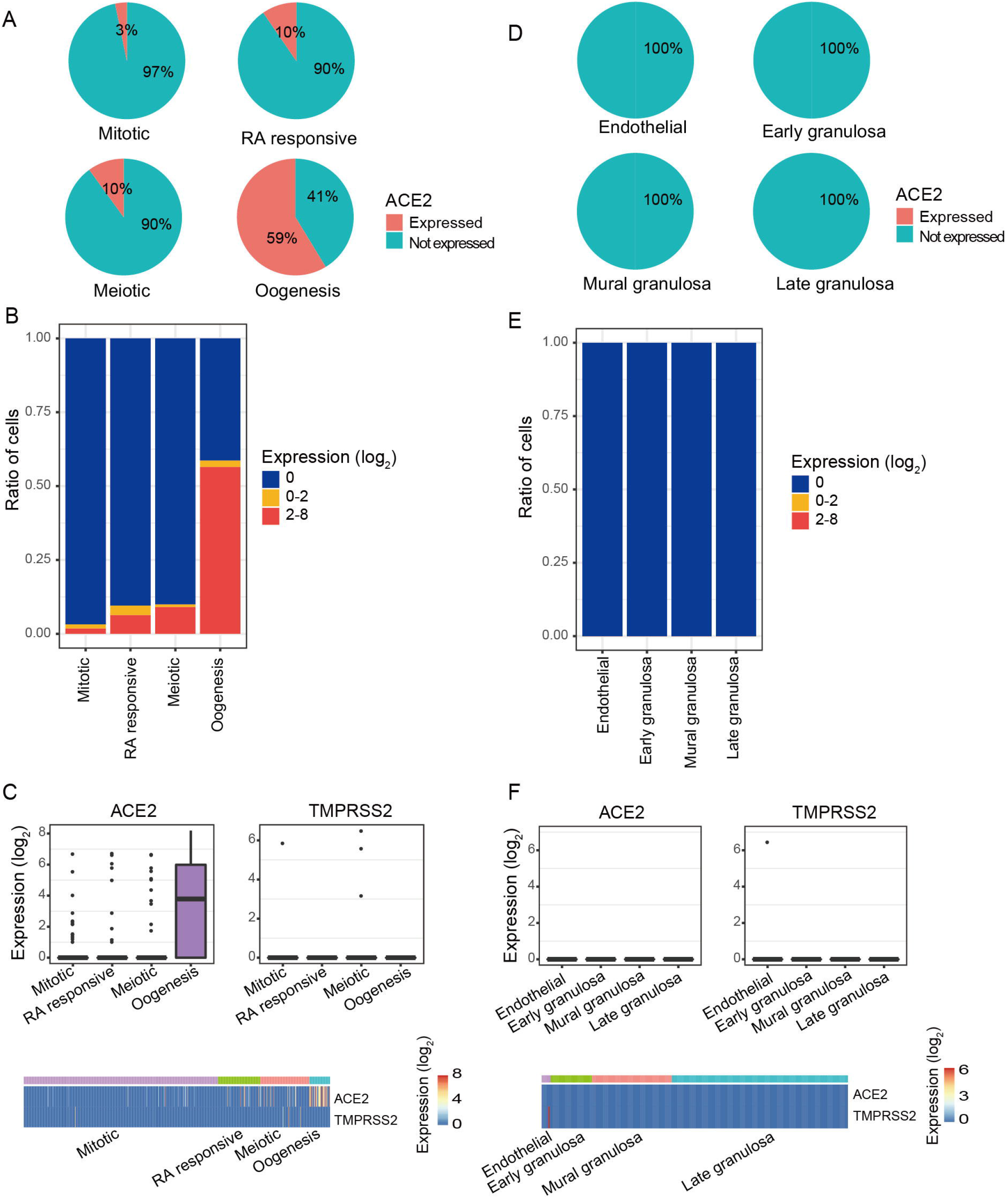
The *ACE2* expression pattern in female PGCs and their surrounding somatic cells. A) Pie charts showed the ratio of *ACE2* expressed PGCs in each developmental stage. Mitotic, RA responsive, Meiotic, and Oogenesis represent four sequentially developmental stages of female PGCs. B) Ratios of different *ACE2* expression level of PGCs in each developmental stage. C) *ACE2* and *TMPRSS2* expression level of PGCs. Top: boxplots of *ACE2* and *TMPRSS2* expression level in each developmental stage. Bottom: heatmaps of *ACE2* and *TMPRSS2* expression level in each cell. D) Pie charts showed the ratio of *ACE2* expressed PGC surrounding somatic cells. Endothelial: somatic cells surrounding the PGC in Mitotic stage; Early granulosa: somatic cells surrounding the PGC in RA responsive stage; Mural granulosa: somatic cells surrounding the PGC in Meiotic stage; Late granulosa: somatic cells surrounding the PGC in Oogenesis stage. E) Ratios of different *ACE2* expression level of PGC surrounding somatic cells. F) *ACE2* and *TMPRSS2* expression level of PGC surrounding somatic cells. Top: boxplots of *ACE2* and *TMPRSS2* expression level in PGCs surrounding somatic cells. Bottom: heatmaps of *ACE2* and *TMPRSS2* expression level in each cell.

### 2.2 Oocytes and cumulus cells displayed high ACE2 expression

Female PGCs develop into immature oocytes in adult women ovary and the immature oocytes develop into mature oocytes through the primordial, primary, secondary, antral, and preovulatory stages (Zhang et al., 2018). To understand the developmental-stage-specific expression pattern of *ACE2* in oocytes and surrounding cumulus cells, we analyzed the *ACE2* expression in folliculogenesis. We firstly calculated the proportion of *ACE2* expressed oocytes in different developmental stages during folliculogenesis. The results showed that *ACE2* were expressed in all five stages (primordial, primary, secondary, antral, and preovulatory follicles) with the least proportion of 52.9% *ACE2*-positive cells in primordial follicles and 100% *ACE2*-positive oocytes from antral to preovulatory stage (Fig 2A, 2B), reminding that oocytes during folliculogenesis may possibly be susceptible to virus infections. Moreover, the ratio of *ACE2*-positive cells was sharply up-regulated in primary follicles, indicating that *ACE2* may play a crucial role in this stage (Fig 2A, 2B). Intriguingly, *TMPRSS2* were also low-expressed while *CTSB* and *CTSL* were expressed in oocytes (Fig. 2C, Fig.S2A), suggesting that CatB/L may also be primed for SARS-CoV-2 invasion. Taken together, oocytes during follicular development may be at high risk of SARS-CoV-2 infection, especially in antral and preovulatory stage.

**Figure 2.**
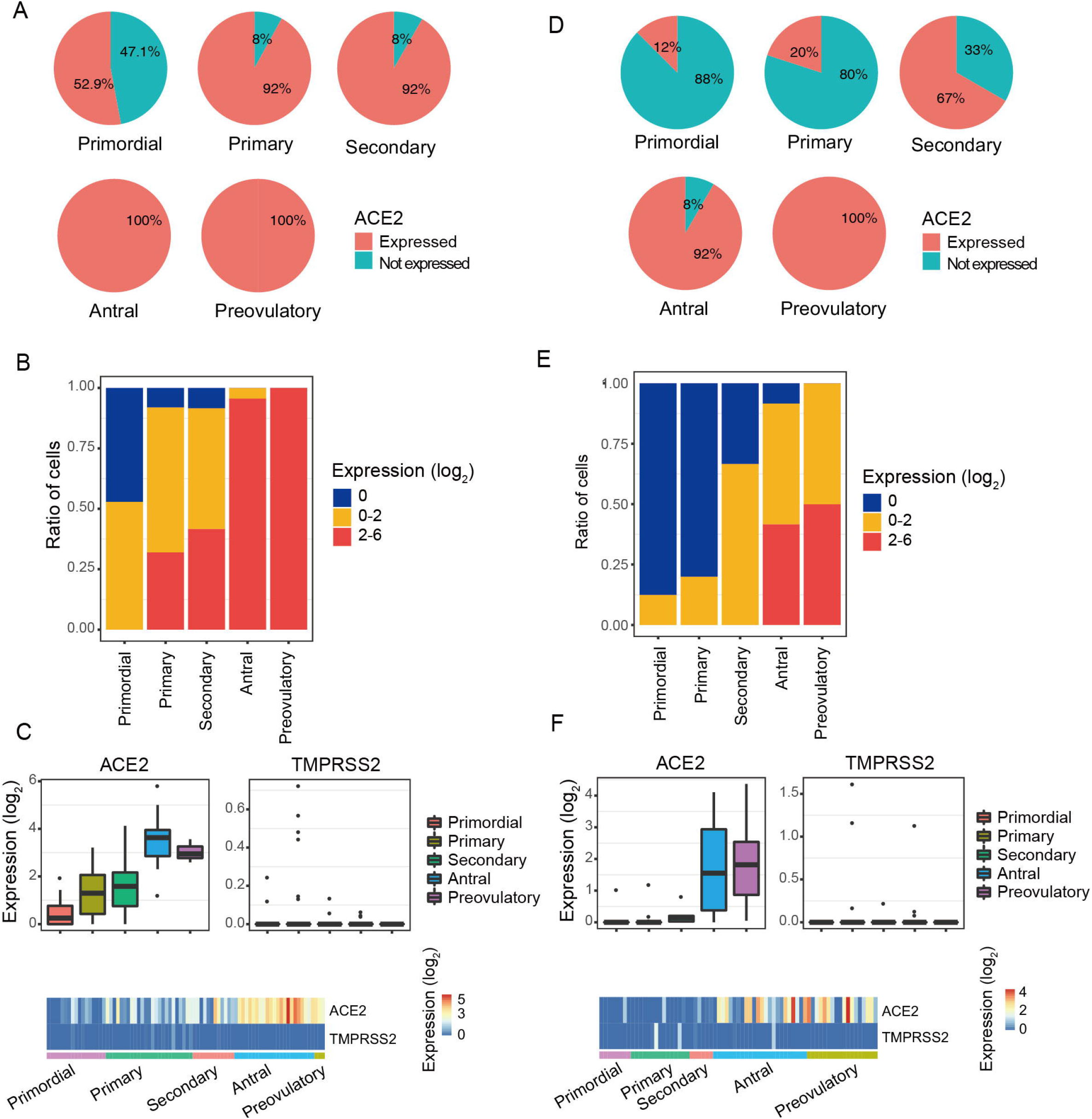
The *ACE2* expression pattern in COC during follicle development. A) Pie charts showed the ratio of *ACE2* expressed oocytes in COC of each developmental stage. Primordial, Primary, Secondary, Antral, and Preovulatory represent the five sequentially developmental stages of oocytes. B) Ratios of different *ACE2* expression level of oocytes in COC of each developmental stage. C) *ACE2* and *TMPRSS2* expression level of oocytes in COC. Top: boxplots of *ACE2* and *TMPRSS2* expression level in each developmental stage. Bottom: heatmaps of *ACE2* and *TMPRSS2* expression level in each cell. D) Pie charts showed the ratio of *ACE2* expressed cumulus cells in COC of each developmental stage. Primordial, Primary, Secondary, Antral, and Preovulatory represent the five sequentially developmental stages of oocyte surrounding granulosa cells. E) Ratios of different *ACE2* expression level of cumulus cells in COC of each developmental stage. F) *ACE2* and *TMPRSS2* expression level of cumulus cells in COC. Top: boxplots of *ACE2* and *TMPRSS2* expression level in each developmental stage. Bottom: heatmaps of *ACE2* and *TMPRSS2* expression level in each cell.

To further understand the effects of *ACE2* on folliculogenesis, we performed Gene Ontology analysis between *ACE2*-positive and *ACE2*-negative oocytes in primary stage. We found that *ACE2* was involved in the regulation of autophagy and in key metabolic process, such as dephosphorylation, glycosylation, and amino acid transport (Fig.S3C).

Meanwhile, we also tracked genes synchronously expressed with *ACE2* in folliculogenesis. Interestingly, we found that *USP13* had the same pattern with *ACE2* during follicular development (Fig.S3A). Likewise, GO analysis in down-regulated genes between *ACE2*-positive and *ACE2*-negative oocytes in primary stage also showed that *ACE2* was involved in cilium movement (Fig.S3C). Together, *ACE2* may be involved in metabolic process, regulation of autophagy and cilium movement during follicular activation and ovulation.

Cumulus cells were neighboring cells of oocytes and may facilitate oocyte maturation (Chang et al., 2016). *ACE2* were expressed in cumulus cells of all five developmental stages, including primordial (12% *ACE2*-positive cells), primary (20% *ACE2*-positive cells), secondary (67% *ACE2*-positive cells), antral (92% *ACE2*-positive cells) and preovulatory (100% *ACE2*-positive cells) stages (Fig 2D, 2E), indicating that cumulus cells during folliculogenesis were at high risk of SARS-CoV-2 infection. To further confirm the SARS-CoV-2 entry, we also compared *TMPRSS2* and *ACE2* expression in cumulus cells. The results showed that there were hardly *TMPRSS2* positive cells in *ACE2* positive cells (Fig 2F). Interestingly, *CTSB* and *CTSL* were expressed in all stages during folliculogenesis (Fig.S2B) which meant that CatB/L may prime SARS-CoV-2 for cell entry. Together, cumulus cells during folliculogenesis may be at high risk for SARS-CoV-2 infection. Collectively, *ACE2* were high expressed in COC and SARS-CoV-2 may use CatB/L for priming.

To better study *ACE2* function in cumulus cells during follicle activation, we also performed Gene Ontology analysis in primary stage. The results showed that up-regulated genes in *ACE2*-positive cumulus cells were involved in the pattern recognition receptor signaling pathway, nucleobase-containing compound transport (Fig.S3D). Down-regulated genes in *ACE2* positive cumulus cells was involved in cilium assembly (Fig.S3D). Interestingly, down-regulated genes in *ACE2* positive oocytes were also related to cilium movement, suggesting that *ACE2* may have influences on cilium movement and cumulus cells may help with this process. Together, these results indicated that cumulus cells may help with the receptor recognition, compound transport and cilium movements during oocyte maturation.

## 3. Discussion

Organs with *ACE2* expression may exhibit potential risk of SARS-CoV-2 infection and *ACE2* bound to SARS-CoV-2 S ectodomain presented 10-to 20-fold higher affinity than *ACE2* binding to SARS associated coronavirus (Wrapp et al., 2020). The SARS-Cov-2 positive detection results in female ovary reminded the infectious risk in female reproduction (W, 2020). However, the developmental-stage-specific and cell-type-specific risk in fetal primordial germ cells (PGCs) and adult women ovary still needs investigation. In this study, we used scRNA-seq datasets from PGCs and COCs to systematically profile *ACE2* expression pattern in female ovary.

Our study first revealed that PGCs displayed high *ACE2* expression in all stages with 59% *ACE2* positive cells in oogenesis phase, indicating the high risk of SARS-CoV-2 susceptibility in PGCs. And SARS-CoV-2 may use CatB/L to prime cell entry. However, whether SARS-CoV-2 can pass through the maternal-fetal barrier (Delorme-Axford et al., 2014) need more research.

We also found that *ACE2* were expressed in oocytes during folliculogenesis and all oocytes in antral and preovulatory stages were identified to express *ACE2*. *ACE2* were sharply up-regulated at the time of oocyte maturation in a rainbow trout study (Bobe et al., 2006). We also found that *ACE2* were expressed in cumulus cells especially in preovulatory stage (100% *ACE2*-positive cells). In addition, *ACE2* were hardly detected in mural granulosa cells. It was consistent with that cumulus cells, not mural granulosa cells, in human preovulatory follicles showed high *ACE2* expression after 36 hours human chorionic gonadotropin administration (Grøndahl et al., 2012). Furthermore, our study found that CatB/L, rather than TMPRSS2, was identified to prime for SARS-CoV-2 entry in COC. Besides, we found that the proportion of *ACE2*-positive cells was sharply up-regulated in oocytes at primary stage and *ACE2* may be related to metabolic process and regulation of autophagy. Autophagy involved USP13 (Xie et al., 2020) is synchronously expressed with *ACE2* during folliculogenesis. In addition, cumulus cells were also found to be related to receptor recognition and compound transport. These reminded us that *ACE2* changes may have influences on oocyte maturation.

Collectively, our study highlighted that the potential risk of SARS-CoV-2 infection in fetal PGCs and adult COCs with high *ACE2* expression. CatB/L may be used to prime SARS-CoV-2 infection. More experimental research and clinical observation were necessary to provide more robust evidence.

## 4. Materials and Methods

We collected the published scRNA-seq datasets from different scales in ovary, including female PGCs (GSE86146) and COC (GSE107746). The data was processed according to the description in corresponding studies of each dataset (Li et al., 2017; Zhang et al., 2018). The expression level was log transformed in this analysis. The cells with *ACE2* expression>0 were defined as *ACE2*-postive cells. According to the accepted consensus that lung AT2 cells are susceptible to SARS-CoV-2, the expression level of *ACE2* in AT2 cells were used as a reference. Cell types with the *ACE2*-positive proportion >1% were regarded as high risk. Student t-test were used to identify differentially expressed genes between *ACE2*-positive and *ACE2*-negtive cells in primary stage in COC. Gene Ontology analysis of the differentially expressed genes was performed using metascape (Zhou et al., 2019).

## Abbreviations

SARS-CoV-2: severe acute respiratory coronavirus 2
SARS: Severe acute respiratory syndrome
PGCs: primordial germ cells
ACE2: angiotensin-converting enzyme2
CatB/L: cathepsin B and L
scRNA-seq: single-cell RNA-sequencing
COC: cumulus-oocyte complex.

## Acknowledgements

This work was supported by National Key Research and Development Program (2019YFA0801400, 2018YFC1004000, 2017YFA0103801) and National Natural Science Foundation of China (81521002, 81730038).

## Data availability

The datasets supporting the conclusions of this article are included in these published articles (Li et al., 2017; Zhang et al., 2018).

## Compliance and ethics

The authors declare that they have no competing interests.

---

**Fig.S1.**
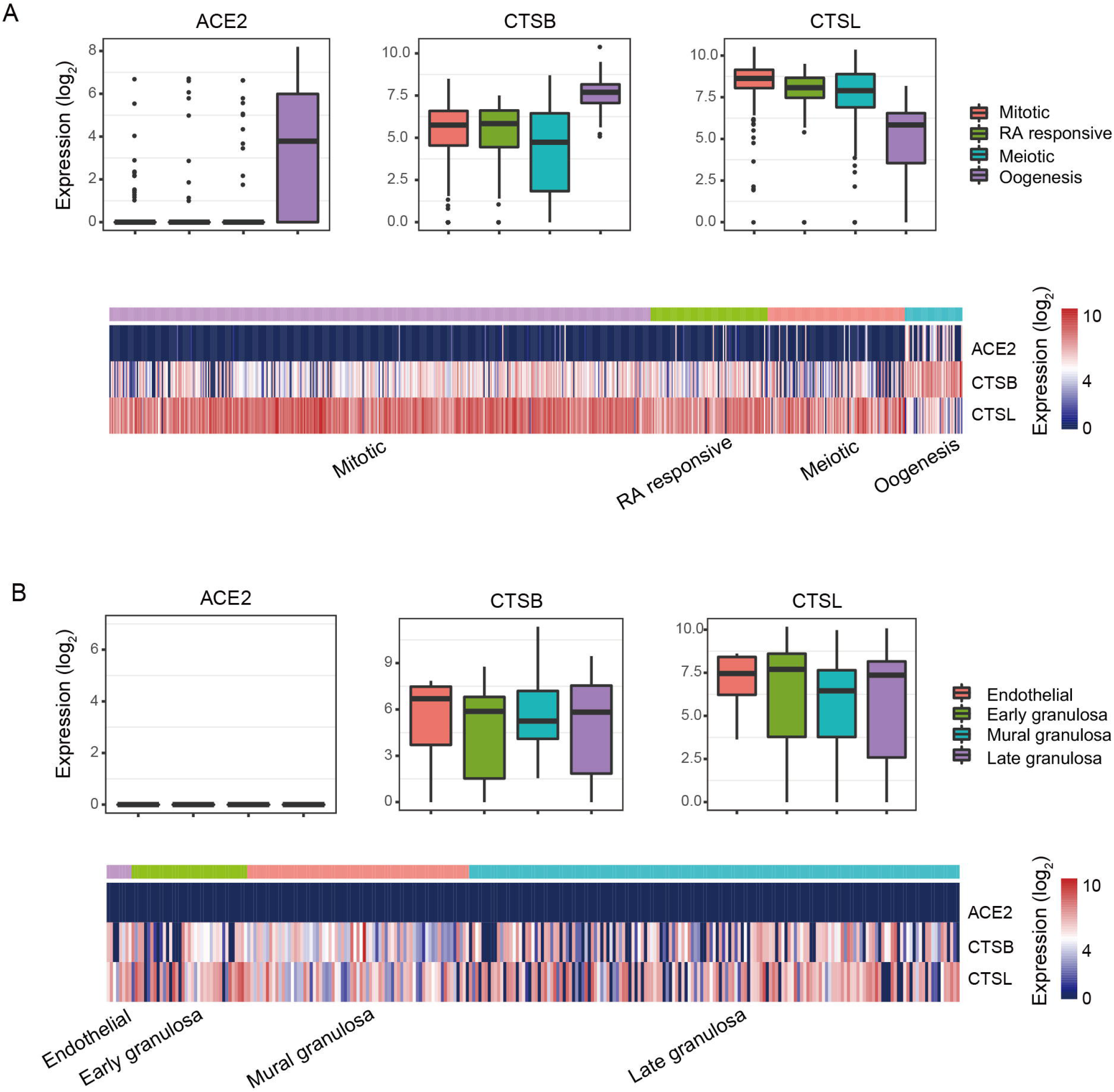
*ACE2*, CTSB and CTSL expression in female embryo PGCs. A) *ACE2*, *CTSB* and *CTSL* expression level in PGCs. Top: boxplots of *ACE2*, *CTSB* and *CTSL* expression level in each developmental stage. Bottom: heatmaps of *ACE2*, *CTSB* and *CTSL* expression level in each cell. B) *ACE2*, *CTSB* and *CTSL* expression level in PGCs surrounding somatic cells. Top: boxplots of *ACE2*, *CTSB* and *CTSL* expression level in PGCs surrounding somatic cells. Bottom: heatmaps of *ACE2*, *CTSB* and *CTSL* expression level in each cell.

**Fig.S2.**
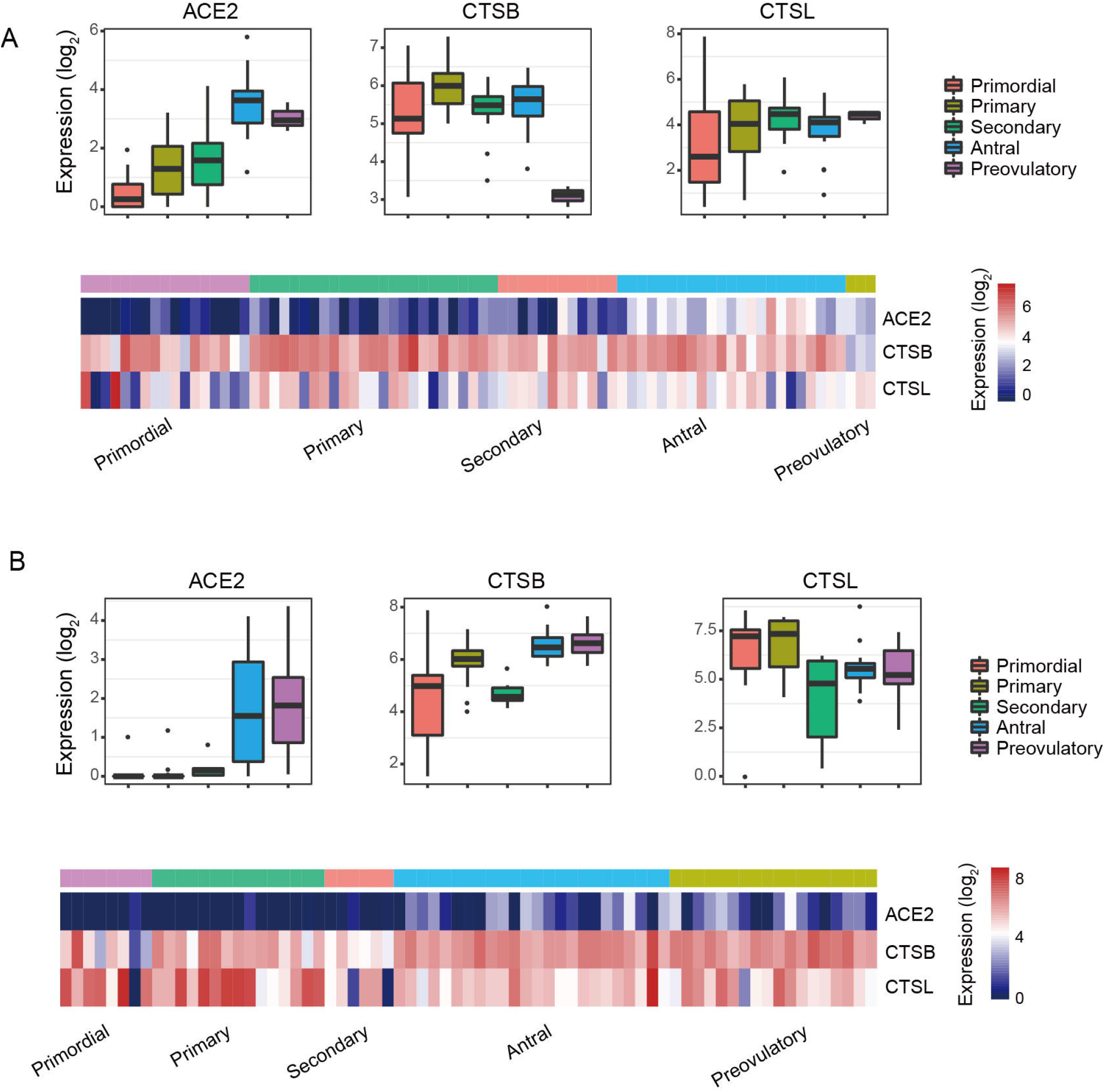
*ACE2*, *CTSB* and *CTSL* expression in human COC during follicle development. A) *ACE2*, *CTSB* and *CTSL* expression level in oocytes. Top: boxplots of *ACE2*, *CTSB* and *CTSL* expression level in each developmental stage. Bottom: heatmaps of *ACE2*, *CTSB* and *CTSL* expression level in each cell. B) *ACE2*, *CTSB* and *CTSL* expression level in surrounding cumulus cells. Top: boxplots of *ACE2*, *CTSB* and *CTSL* expression level in each developmental stage. Bottom: heatmaps of *ACE2*, *CTSB* and *CTSL* expression level in each cell.

**Fig.S3.**
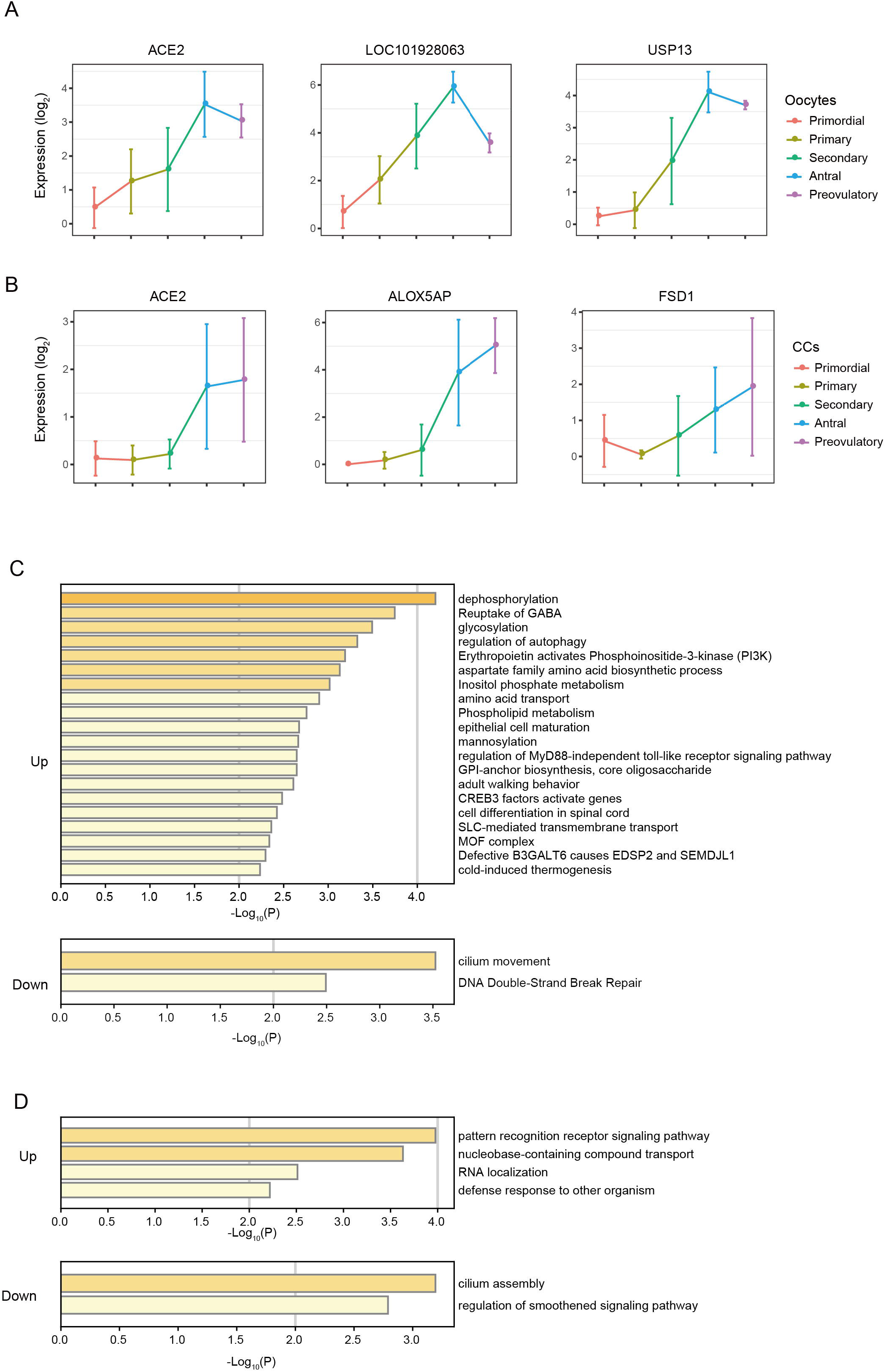
Gene Ontology analysis of genes between *ACE2*-positive and *ACE2*-negative cells. A) Genes synchronously expressed with *ACE2* in oocytes. B) Genes synchronously expressed with *ACE2* in cumulus cell. C) Gene ontology terms of genes in oocytes between *ACE2*-positive and *ACE2*-negative cells in primary stage. Top: gene ontology terms of up-regulated genes in oocytes. Bottom: gene ontology terms of down-regulated genes in oocytes. D) Gene ontology terms of genes in cumulus cells between *ACE2*-positive and *ACE2*-negative cells in primary stage. Top: gene ontology terms of up-regulated genes in cumulus cells. Bottom: gene ontology terms of down-regulated genes in cumulus cells.

